# Development of a novel microphysiological system for neurotoxicity prediction using human iPSC-derived neurons with morphological deep learning

**DOI:** 10.1101/2024.06.08.598041

**Authors:** Xiaobo Han, Naoki Matsuda, Makoto Yamanaka, Ikuro Suzuki

**Affiliations:** Department of Electronics, Graduate School of Engineering, Tohoku Institute of Technology, 35-1 Yagiyama Kasumicho, Taihaku-ku, Sendai, Miyagi, 982-8577, Japan; Ushio Inc

## Abstract

Microphysiological system (MPS) is an in vitro culture technology that reproduces the physiological microenvironment and functionality of humans, and is expected to be applied for drug screening. In this study, we developed a MPS for structured culture of human iPSC-derived neurons, then predict drug-induced neurotoxicity by morphological deep learning.

For human iPSC-derived cortical neurons, after administration of three different amyloid β (Aβ) peptides, neurotoxicity effects were evaluated by a deep learning image analysis, training two artificial intelligence (AI) models on neurites and PSD-95 images. The combined results indicated that Aβ 1-42 and 1-40, but not 1-28, induced synaptic degeneration, which is close to clinical reports. For human iPSC-derived sensory neurons, after administration of representative CIPN-related anti-cancer drugs, the toxic effects on soma and axons were evaluated by AI model using neurites images. Significant toxicity was detected in positive drugs and could be classified by different effects on soma or axon, suggesting that the current method provides an effective evaluation of chemotherapy-induced peripheral neuropathy. Taken together, it suggests that the present MPS combined with morphological deep learning is a useful platform for in vitro neurotoxicity assessment.

## Introduction

Novel therapeutics are necessary to treat neurological disorders such as Alzheimer’s disease (AD), Parkinson’s disease, multiple sclerosis and amyotrophic lateral sclerosis (ALS). Globally, in 2016, neurological disorders were the leading cause of disability-adjusted life years (276 million) and second leading cause of deaths (9 million) [1]. Notably, 15% of children in the US ages 3 to 17 yr were affected by neurodevelopmental disorders [2]. However, there are not enough therapeutic options for neurological disorders, as most approaches merely address symptom alleviation [3,4]. To address the need for streamlining within the drug development, more physiologically relevant models are needed to screen potential candidate compounds in nascent testing stages [5,6]. The overall goal should be to rule out cytotoxic drugs and drugs without human efficacy earlier, thus reducing time and money spent on determining appropriate candidates. Understandings in neuronal functionality that can be studied using discrete, testable microphysiological systems (MPS) are imperative to the development of therapeutics that move more towards regenerative therapies.

The advent of stem cell technologies and bioengineering in cell culture have led to the recent development of MPS which are expected to capture the human-like physiologies of tissues and organs in vitro, i.e., the structures and physiochemical factors of the tissues [7–13]. The neural MPS normally should be constructed with the proper cell sources, materials, and fabrication methods, to recapitulate some organ-level functionality without requiring in situ or in vivo methods. In the present study, we developed a MPS device for culturing human iPS cell-derived neurons. This MPS device provides structured neural culture that separates neurites grow-out into the microfluidic channels. Human iPSC-derived cortical neurons or sensory neurons were cultured in the MPS device. Then the morphological changes in neurons under drug administrations were analyzed by a deep learning method. After training AI models with images obtained from the MPS device, drug-induced neurotoxicity was detected for each compound.

## Material and methods

### MPS device fabrication

Ushio Inc. prepared the MPS device as previously described [14]. Briefly, the vacuum ultraviolet (VUV) photobonding from an excimer light at a 172-nm wavelength was used to generate the functional groups (i.e., hydroxy and carboxyl groups) for assembling two COP material layers directly with heat treatment. One MPS device is comprised of four individual microfluidic cell culture channels (Figure 1A-a). The middle narrow slot part of the channel is 1000 µm in width, 165 µm in length, and 40 µm in height, with an open-top channel (1000 µm in width, 6 mm in length) and two circular holes (2 mm in diameter) at both ends which open to a rectangular medium reservoir (15 mm in width, 8 mm in length, and 5 mm in height). The maximum volume in each channel containing reservoirs is 1 mL. COP material (Zeonex 690R, Zeon, Tokyo, JAPAN) was injected to the two molds individually. The components were irradiated with VUV from an excimer lamp (172 nm; Ushio Inc., Tokyo, JAPAN) at 25°C after taking the structured COP components from the molds. The component surfaces were assembled using a heat press at less than 132°C. Finally, ethylene oxide gas (Japan Gas Co. Ltd., Kanagawa, JAPAN) was used for device decontamination.

**Figure 1.**
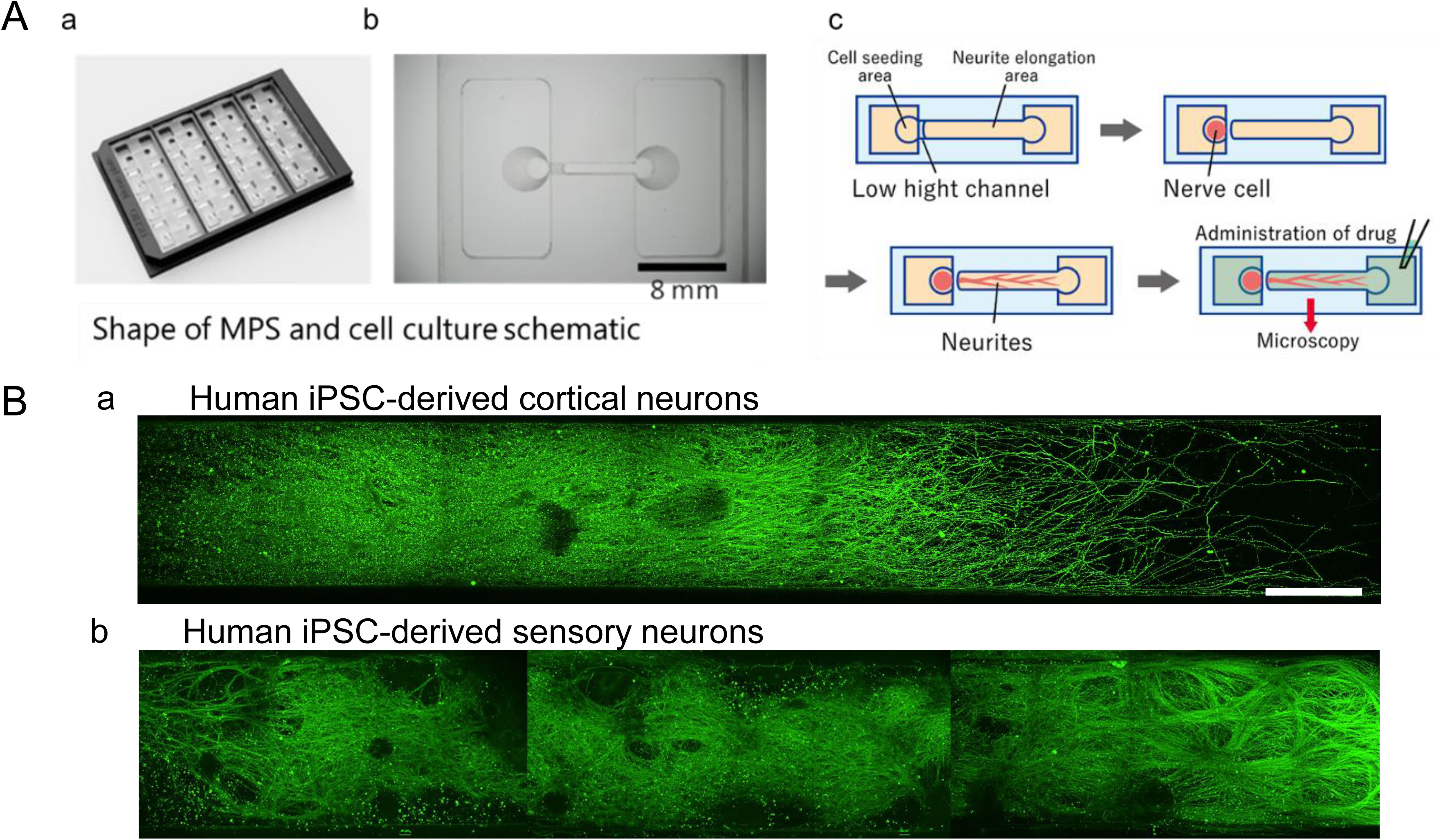
Shape of the microphysiological system (MPS) and neurites elongation images. (A) MPS with (a) an overview. (b) Magnified view of a single channel. Scale bar = 8 mm. (c) Schematic of cultivation. (B) Representative immunofluorescence images of neurites elongation area for (a) human iPSC-derived cortical neurons and (b) human iPSC-derived sensory neurons. Scale bar = 500 µm.

### Cell culture

Before seeding human iPSC-derived cortical neurons, the surface of MPS device was coated sequentially by Cellmatrix collagen type I-C solution (637-00773, Nitta Gelatin), Poly-D-lysine solution (P7405, Sigma-Aldrich), and iMatrix 511 laminin solution (892019, Matrixome), each for 1 h. Cryopreserved human iPSC-derived cortical neurons (XCL-1 Neurons, XCell Science) were thawed and suspended in Neuron Medium (XCS-NM-001-M100-1P, XCell Science). For dispersed culture, approximately 5.0 × 10^4^ cells in 15 µL neuron medium were seeded directly into the seeding chamber at one side of the MPS device. After 30 min, totally 600 µL of neural maturation basal medium (NM-001-BM100, XCell Science Inc., USA) supplemented with neuron maturation supplement A (NM-001-SA100, XCell Science Inc., USA) and 100 U/mL penicillin/streptomycin (168– 23191, Wako) was applied into the whole MPS device. After one week, the medium was replaced with 600 µL of BrainPhys neuronal medium containing SM1 neuronal supplement (ST-05792, STEMCELL technologies). Following this, half the volume of the medium was replaced twice per week. After 6 weeks in culture, three typical amyloid β (Aβ) peptides were administered to the cultures at two different concentrations each: Aβ 1-28 at 0.1 µM and 1 µM, Aβ 1-40 at 0.1 µM and 1 µM, Aβ 1-42 at 0.1 µM and 1 µM. As a comparative compound, KT-430 at 10 µM and 100 µM, a low molecular compound derived from isoproterenol that inhibits aggregation of tau protein, was administered. DMSO (0.1%) was added as the negative control drug to the cultures. The drug exposure lasted for 24 h at 37 ℃.

Before seeding human iPSC-derived sensory neurons, the surface of MPS device was coated sequentially by Cellmatrix collagen type I-C solution (637-00773, Nitta Gelatin), Poly-D-lysine solution (P7405, Sigma-Aldrich), and SureBond-XF coating solution (AXOL Bioscience, ax0053), each for 1 h. All plating and maintenance media were obtained from Axol’s kit (ax0157, Axol Bioscience). Cryopreserved human iPSC-derived Sensory Neuron Progenitors (ax0055, Axol Bioscience) were thawed and suspended in the plating medium. For dispersed culture, approximately 5.0 × 10^4^ cells in 15 µL neuron medium were seeded directly into the seeding chamber at one side of the MPS device. After 30min, totally 600 µL of plating medium was applied into the whole MPS device. The total volume of medium was replaced with maintenance medium containing growth factor (Sensory maturation maximazer supplement, GDNF: 25ng/mL, ax139855, NGF: 25ng/mL, ax139789, BDNF: 10ng/mL, ax139800, NT-3: 10ng/mL, ax139811) and maximizer (sensory maintenance medium) after 24 h. Following this, half of the medium was replaced every 3 days. After 3 weeks (18∼20 days) in culture, three typical anticancer drugs were administered to the cultures at two different concentrations each: paclitaxel at 0.1 µM and 1 µM, vincristine at 0.003 µM and 0.03 µM, oxaliplatin at 10 µM and 100 µM. As comparative compounds, bortezomib at 0.01 µM, a proteasome inhibitor, and suramin at 10 µM and 100 µM, antiparasitic drugs that have antineoplastic effects and cause myelin damage, were administered. DMSO (0.1%) was added as the negative control drug to the cultures. The drug exposure lasted for 24 h at 37 ℃.

### Immunocytochemistry

Sample cultures were fixed with 4% paraformaldehyde in PBS on ice (4°C) for 10 min. Fixed cells were incubated with 0.2% Triton-X-100 in PBS for 5 min, then with preblock buffer (0.05% Triton-X and 5% FBS in PBS) at 4°C for 1 h, and finally with preblock buffer containing specific primary antibodies at 4°C for 24 h. Primary antibodies are listed as: mouse anti-β-tubulin III (1:1000, T8578, Sigma–Aldrich), rabbit anti PSD-95 (1:200, ab18258, Abcam) for human iPSC-derived cortical neuron; mouse anti-β-tubulin III for human iPSC-derived sensory neuron. Samples were then incubated with the secondary antibodies (i.e., anti-mouse 488 Alexa Fluor (1:1000 in preblock buffer, A-11001, Invitrogen), and anti-rabbit 546 Alexa Fluor (1:1000 in preblock buffer, A-11035, Invitrogen)) for 1 h at room temperature. Stained cultures were washed twice with preblock buffer and rinsed twice with PBS. An inverted microscope with confocal imaging system (AX/R, Nikon) was used to visualize immunolabeling for local images and whole microchannel-length images. The ImageJ software (NIH) was used to adjust the image intensity.

### Deep learning for image analysis and toxicity positive prediction

Image treatment and AI creating were performed in a similar method as described before [14]. Briefly, for human iPSC-derived cortical neurons, total immunofluorescence images of neurites and PSD-95 from cell seeding areas and neurite elongation areas were firstly segmented into 576 x 576-pixel images, then used as image datasets for training and test of the neurotoxicity prediction AI analysis. The percentage of segmented images that were determined to be positive was calculated as the toxicity probability. For human iPSC-derived sensory neurons, immunofluorescence images of neurites from neurite elongation areas were segmented into 576 x 576-pixel images and used for AI analysis. Vincristine images were trained as neurite toxicity positive, and oxaliplatin as cytotoxicity positive, then all segmented images were tested and the percentage that were determined to be cytotoxicity/neurite toxicity positive was calculated as the toxicity probability.

### Statistical analysis

The one-way ANOVA was used to perform multiple group comparisons followed by Dunnett’s test or Holm’s test, which were used to calculate the significant difference between each compound at different concentration.

## Results

### Morphological presentation of human iPSC-derived cortical neurons in the MPS device

Human iPSC-derived sensory neurons were exposed to various Aβ peptides after 6 weeks of culture and, subsequently, 24 h post exposure, immunostained images were captured. Tested compounds included Aβ 1-40 and Aβ 1-42, with key relationship to AD, as well as Aβ 1-28, with less relationship to AD. KT-430, which is associated with microtubules and Tau protein aggregation was chosen for comparison, and DMSO was utilized as negative control. Figure 2 shows the local immunofluorescence images of cultured human iPSC-derived cortical neurons at 6 weeks in the MPS device after drug administrations. Morphological alterations of either neurites or PSD-95, a post-synaptic marker, in response to the compounds were evaluated. Under negative control DMSO, neurites elongation was observed together with the expression of PSD-95, which indicated the well formation of neural synaptic network. Compared to DMSO, cell aggregation was observed at some points under administration of Aβ 1-28. But neurites degeneration was not observed and PSD-95 expression was similar to that under DMSO. More cell aggregation was observed in samples under administration of Aβ 1-40 and Aβ 1-42, especially under high concentrations. Increasing of PSD-95 block was observed in samples under administration of Aβ 1-40, and was more severe under Aβ 1-42 even with obviously fragments formation at high concentration. Under administration of KT-430, neurites degeneration was observed clearly.

**Figure 2.**
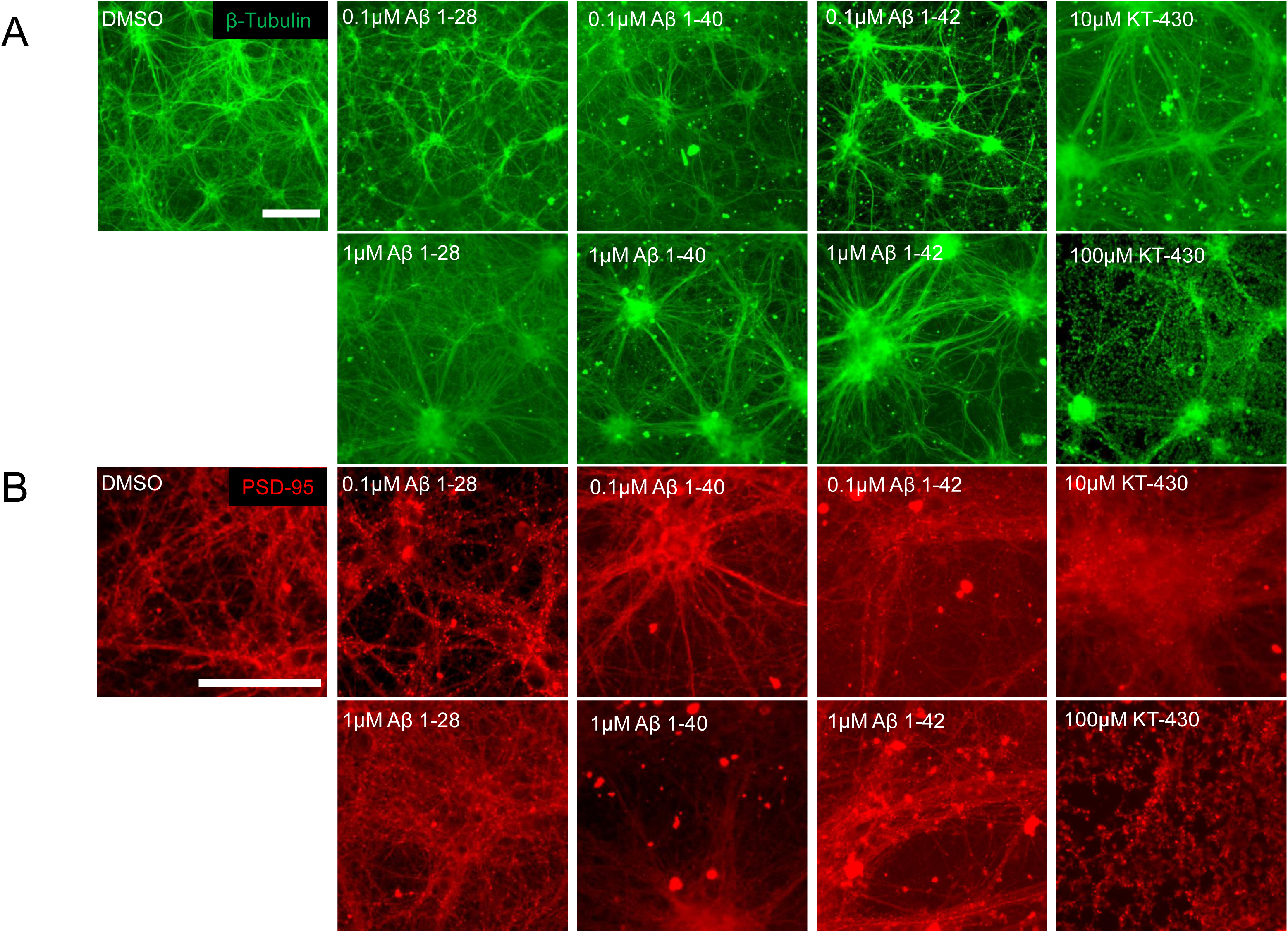
Representative immunofluorescence image samples of human iPSC-derived cortical neurons in the MPS device after drug administration with staining of (A) β-Tubulin, and (B) PSD-95. Scale bar = 50 μm.

### Neurotoxicity prediction for human iPSC-derived cortical neuron using morphological deep learning

Two AI models for neurotoxicity prediction were created and trained with either neurites image datasets or PSD-95 image datasets as described above. Figure 3C showed the prediction rate of toxicity positivity for each compound with neurites image datasets. The negative control compound, DMSO showed a low toxicity positive rate. Aβ 1-40 and KT-430 at both low and high concentrations showed high toxicity positive rates with significant difference compared to DMSO. Figure showed the prediction rate of toxicity positivity for each compound with PSD-95 image datasets. Similarly, DMOS showed a low toxicity positive rate. Aβ 1-28 at high concentration, Aβ 1-40 at low and high concentration, Aβ 1-42 at high concentration, and KT-430 at low and high concentration showed high toxicity positive rates with significant difference compared to DMSO.

**Figure 3.**
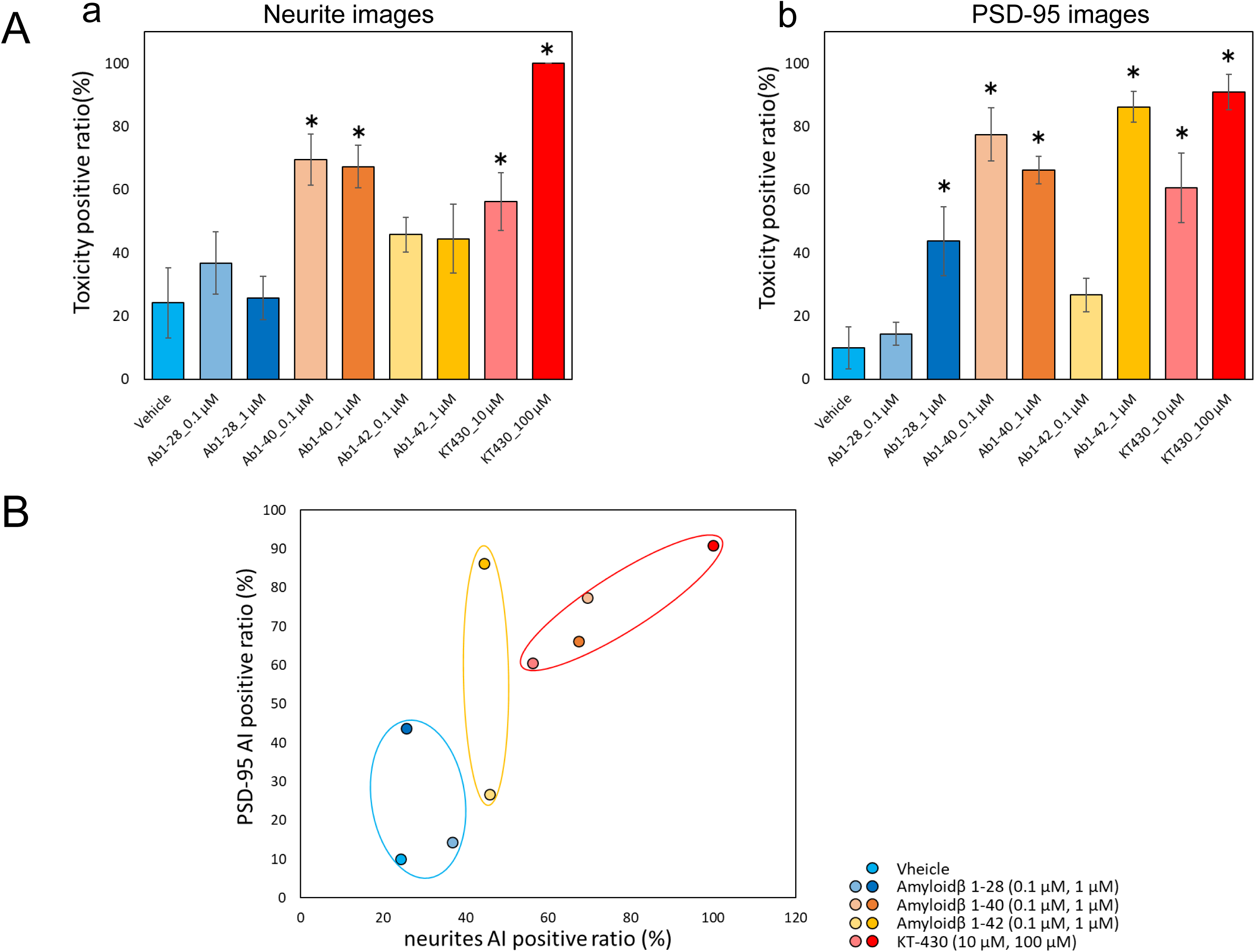
Predicted toxicity positive rates of various compounds on human iPSC-derived cortical neuron by AI analysis. (A) The neurotoxicity positive rate predicted by AI analysis for each drug based on either (a) neurites image datasets, or (b) PSD-95 image datasets. Data were expressed by means ∓ standard errors. ANOVA was used to perform statistical analysis followed by post hoc Dunnet’s test or Holm’s test, * p < 0.05 vs. vehicle. (B) Classification of the mechanisms of each compound based on the combined results of two AI models. The toxicity probability by PSD-95 AI was taken as the vertical axis and by neurites AI as the horizontal axis.

To classify the MoA of compounds on neurons, we utilized the toxicity positive results from two above AI models. Specifically, the toxicity positive rate with neurites images was taken as the vertical axis and with PSD-95 images as the horizontal axis. The outcomes for each compound were plotted to examine their distributions (Figure 4). The graphical representation indicated that DMSO and Aβ 1-28 clustered near the origin. Meanwhile, Aβ 1-42 exhibited a predominant shift in the y-axis direction, indicating its pronounced effect on PSD-95. Aβ 1-40 and KT-430 showed a similar tendency that shifted in the upper-right quadrant.

**Figure 4.**
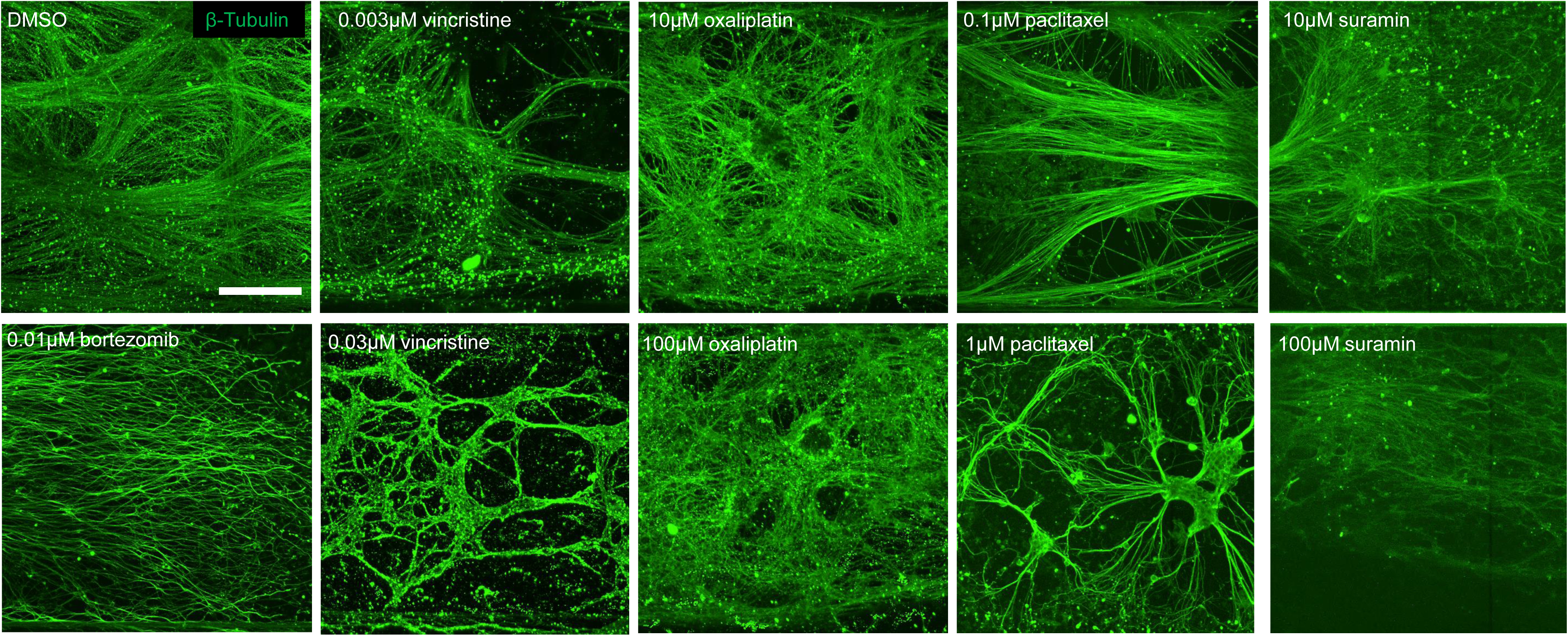
Representative immunofluorescence image samples of human iPSC-derived sensory neurons in the MPS device after drug administration with staining of β-Tubulin. Scale bar = 50 μm.

**Figure 5.**
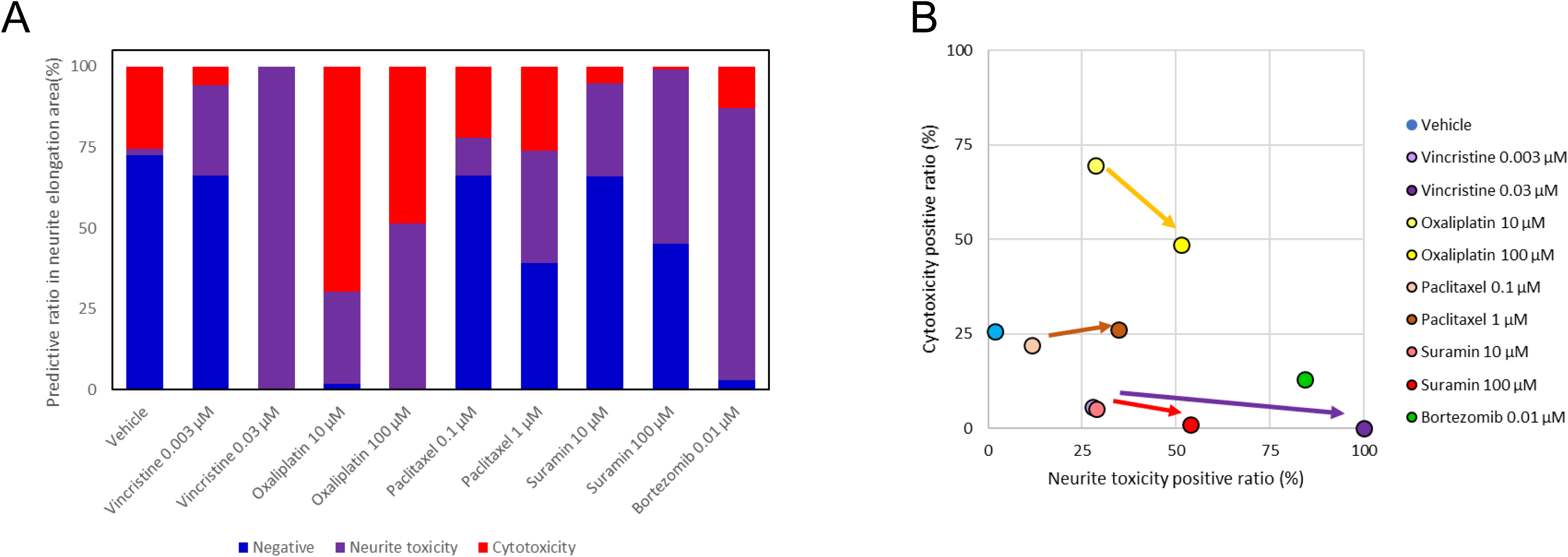
Predicted toxicity positive rates of various compounds on human iPSC-derived sensory neuron by AI analysis. (A) The neurotoxicity positive rate predicted by AI analysis for each drug based on neurites image datasets. Data were expressed by the percentages of negative rate (in blue), neurite toxicity positive rate (in purple), and cytotoxicity positive rate (in red). (B) Classification of the mechanisms of each compound. The neurite toxicity probability rate was taken as the vertical axis and cytotoxicity probability rate as the horizontal axis.

### Morphological presentation of human iPSC-derived sensory neurons in the MPS device

Human iPSC-derived sensory neurons were exposed to various anticancer drugs after 3 weeks of culture and, subsequently, 24 h post exposure, immunostained images were captured. Tested drugs included paclitaxel and vincristine, known inducers of axonal damage, as well as oxaliplatin, which causes soma damage. DMSO was utilized as negative control. For validation, bortezomib, a proteasome inhibitor associated with CIPN, and suramin, an antiparasitic drug with antineoplastic effects but known to damage myelin, were chosen. Figure 2 shows the local immunofluorescence images of cultured human iPSC-derived sensory neurons at 3 weeks in the MPS device after drug administrations. Morphological alterations in response to the compounds were evaluated within the micro-channel area. Under the negative control DMSO, unidirectional neurites elongation was observed throughout the microchannel. Under administration of 0.003 µM vincristine, axonal degeneration was observed in forms of fragment and hollowing out. Thus phenomenon went more clear under 0.03 µM vincristine. Under administration of oxaliplatin at low and high concentrations, axonal aggregation was observed. A clear depletion in axon was observed under administrations of paclitaxel, suramin, and bortezomib.

### Neurotoxicity prediction for human iPSC-derived sensory neuron using morphological deep learning

In order to predict neurotoxicity and the influence point (i.e., cytotoxicity, or neurite toxicity), an AI was created by training with DMSO image datasets as negative, oxaliplatin images as cytotoxicity, vincristine images as neurite toxicity as described above. And the predictive neurotoxicity positive ratio for each compound was show in Figure 4. DMSO was detected as negative with a high negative ratio. For vincristine, the neurite toxicity positive ration showed a significant increasing and reached maximum at high concentration. Oxaliplatin was predominant detected as cytotoxicity positive, but with an increasing in the neurite toxicity positive ratio at high concentration. A dose-dependent increasing in the neurite toxicity positive ratio was observed under administration of paclitaxel and suramin. Bortezomib was predicted as neurite toxicity with a high positive rate. The outcomes for each compound were plotted to examine their distributions with the cytotoxicity positive ratio as the vertical axis and the neurite toxicity ratio as the horizontal axis. Oxaliplatin exhibited a shift in the y-axis direction, while vincristine primarily moved in the x-axis direction. Paclitaxel shifted in the upper-right quadrant, indicating a simultaneous increase in both axonopathy and cytotoxicity. Suramin also exhibited a dose-dependent shift along the x-axis, similar to that of vincristine. And bortezomib positioned itself close to the vincristine position at high concentration.

## Discussion

Culturing of neuronal and glial cells has been extensively reviewed prior [15,16]. While many of these culture systems focus on non-human cultures, emerging protocols for the rapid generation of human nervous systems from iPSC is of particular interest to MPS development. In this study, we developed a COP-based MPS device which was proven to maintain human iPSC-derived neuron growth without undesired cellular damage. The microchannel structure ensured rapid neurites growth-out with unidirectional elongation. This strategy allows screening applications for human iPSC-derived cortical neuron at 6 weeks of culture, and human iPSC-derived sensory neurons at 3 weeks of culture, which provides potential to investigate human-reliable pathological factors.

Functionality of a neuronal system is paramount for any meaningful data acquisition. Microscopy utilizing fluorescent dyes has allowed for the assessment of physiological activity in many tissue types, including neurons [17,18]. In the present study, we attempted to quantify drug-induced neuron degenerations with microscopic images with the deep learning analysis method. For human iPSC-derived cortical neurons, β-Tubulin III, which stained neurites, and PSD-95, a post-synaptic marker were selected to represent responses in neural synaptic networks. The accumulation of Aβ is wildly reported as a core pathway of AD pathophysiology, while the detailed molecular mechanisms of the pathway and the spatial-temporal dynamics leading to synaptic failure, neurodegeneration, and clinical onset are still under intense investigation [19,20]. In the present study, three different types of Aβ peptides were tested on neurons cultured in the MPS device. By integrating two AI models of neurites or PSD-95, Aβ 1-28 was clustered same to the negative control DMSO. Generally, Aβ 1-28 is at unfolded state to avoid aggregation, and the transformation of the solution structure under certain media condition is probably related to amyloid formation in AD [21], which was different from the media condition in the current study. Meanwhile, Aβ 1-40 and Aβ 1-42 with hydrophobic C-terminal show greater propensity to form plaques in AD [22,23]. And the current deep learning analysis also showed a more directly neurotoxicity positive rate for these two peptides. Especially, the AI model of PSD-95 images indicated a dose dependent increasing of neurotoxicity positive rate in Aβ 1-42. As reported in previous animal studies [24,25], Aβ 1-42 induced rapid suppression in synapse spontaneous activity, which agreed with our current result. Aβ 1-40 showed influence both on neurites and PSD-95, with a similar tendency to KT-430. Axonal degeneration was observed in rodent primary cortical neurons after Aβ 1-40 treating, which is same to the current observation [25]. KT-430 is a low molecular compounds that related to Tau protein aggregation, which has a significant correlation between Aβ 1-40 as reported [26,27]. This could explain our findings, but further experiments on Tau expression are required. Recently, Aβ 1-42/1-40 ratio is listed as a robust measure for detecting amyloid plaques and being utilized to aid in the diagnosis of AD [28–30]. The difference between Aβ 1-40 and 1-42 at structure, function, aggregates formation or other aspects was explored by flexible approaches previously. Our current results also showed a significant difference between these two Aβ peptides, which indicated that the current MPS device provides reliable toxicity prediction.

Anticancer drugs used in cancer treatment could induce chemotherapy-induced peripheral neuropathy (CIPN) as an adverse effect [31]. In the present study, influence of several representative anticancer drugs were detected with human iPSC-derived sensory neurons cultured in the MPS device. The micro-channel structure allowed abundant neurites growth out with unidirectional elongation. After training with image datasets from the micro-channel area, test compounds could be separated as cytotoxicity or neurite toxicity. Previously, our group have reported a similar approach to predict neurotoxicity induced by anti-cancer drugs using rodent primary sensory neurons cultured in the MPS device [14]. Compared to the previous result, the plot map of each compounds showed a same distribution in the current study, which indicates the potential for valid drug screening beyond species gap. However, the morphological of cell bodies were analyzed and trained to AI models in the previous research. Same treatment was not performed in the present study, since the cell bodies of human iPSC-derived sensory neuron were much smaller in size, then difficult to separate individually for deep learning. New approaches of cell culture and image treatment are now under development to enable analysis including cell bodies for a more accurate prediction.

Taken together, a novel COP based MPS device was constructed in the present study. It provides relatively rapid culture timescales for human iPSC-derived cortical neurons and sensory neurons used in the current study, with the micro-channel structure enabled plenty neurites growth out. After morphological deep learning analysis using neurite images,

## Acknowledgement

This study was supported by the grant of Ushio Inc. as a collaborative project.

